# A Bayesian predictive approach for dealing with pseudoreplication

**DOI:** 10.1101/839894

**Authors:** Stanley E. Lazic, Jack R. Mellor, Michael C. Ashby, Marcus R. Munafo

**Affiliations:** Prioris.ai Inc., 459–207 Bank Street, Ottawa, K2P 2N2, Canada; Centre for Synaptic Plasticity, School of Physiology, Pharmacology and Neuroscience, University of Bristol, Bristol, BS8 1TD, UK; MRC Integrative Epidemiology Unit, UK Centre for Tobacco and Alcohol Studies, School of Experimental Psychology, University of Bristol, Bristol, BS8 1TU, UK

## Abstract

Pseudoreplication occurs when the number of measured values or data points exceeds the number of genuine replicates, and when the statistical analysis treats all data points as independent and thus fully contributing to the result. By artificially inflating the sample size, pseudoreplication contributes to irreproducibility, and it is a pervasive problem in biological research. In some fields, more than half of published experiments have pseudoreplication – making it one of the biggest threats to inferential validity. Researchers may be reluctant to use appropriate statistical methods if their hypothesis is about the pseudoreplicates and not the genuine replicates; for example, when an intervention is applied to pregnant female rodents (genuine replicates) but the hypothesis is about the effect on the multiple offspring (pseudoreplicates). We propose using a Bayesian predictive approach, which enables researchers to make valid inferences about biological entities of interest, even if they are pseudoreplicates, and show the benefits of this approach using two *in vivo* data sets.

## Introduction

For nearly a century, statisticians and quantitative biologists have worried about pseudoreplication – also known as the unit-of-anlysis problem.^1^ Hurlbert’s seminal 1984 paper drew attention to the problem of pseudoreplication in the biological sciences, but the concept remains controversial.^2–5^ The requirements for genuine replication are well-defined^6,7^, but how they apply to one’s experiment, or the degree to which the requirements are met can be debatable, as some rely on untestable assumptions. Reasonable disagreement can therefore exist.

The controversy may persist because formulating a scientific question as statistical model can be difficult if one has received little formal training in experimental design or statistics – like many laboratory-based biologists. In addition, for some experiments, the intuitively obvious model may in fact be incorrect. The two main ways of dealing with pseudoreplication are: (1) average the pseudoreplicates to obtain one value per genuine replicate, or (2) use a more sophisticated approach that captures the structure of the data where the pseudoreplicates are nested under the genuine replicates, such as a multilevel/hierarchical model.^8–11^

These recommended methods have lead to a key point of contention. Although statistically appropriate, averaging and multilevel models test a hypothesis at a biological or technical level that differs from researchers’ desired target. For example, a researcher may be interested in how a prenatal intervention affects *offspring’s* behaviour, not how control *litters* differ from treated litters; or how a drug applied to wells in a microtitre plate effects *cellular* morphology, not how control *wells* differ from drug-treated wells; or how a treatment applied to animals affects the number of *synapses* in a brain region, not how the two groups of *animals* differ. Thus, there is a disconnect between what researchers want and what the usual statistical methods provide.

The proposed solution is to cast the problem within a predictive framework. The “predictivist” approach to scientific inference has a long history in statistics. In 1920 Karl Pearson claimed the “fundamental problem of practical statistics” was predicting future observables from past data,^12^ and in 1942, Deming wrote, “Every empirical statement of science is a prediction... There is no scientific interest in any measurement or empirical relationship that does not help to explain what will happen or has happened at another time or another place.”^13^ For Deming, it follows that “It is the statistician’s job to make rational predictions concerning measurements yet to be made.”^13^ But in the early 20th century, statisticians began focusing on parameters in statistical models – estimating their values and making them targets of hypothesis tests.^14,15^ The predictive perspective has since been a minority view in statistics but has developed in parallel to the mainstream’s emphasis on parameters.^14,15^ Fortunately, there is a renewed interest in prediction,^16–20^ perhaps due to the attention that machine learning is receiving.

Emphasising prediction for scientific discovery has many advantages (see Billheimer and Briggs for recent discussions^17,20^), and here we focus on one that addresses the problem of pseudoreplication: we can make statistical inferences about biological entities of interest from multilevel models, that is, about the pseudoreplicates. A second advantage not specific to pseudoreplication is that conclusions are probabilistic statements about observable events, not unobservable parameters, and as Billheimer and Briggs have argued, this can help with the reproducibility crisis.^17,20^ Briggs writes, “Do not speak of parameters; talk of reality, of observables. This alone will go miles toward eliminating the so-called replication crisis.”[17; p. xii]

Focusing on parameters makes more sense when a true but unknown value exists, such as the speed of light, the electrical conductivity of copper, or the boiling point of ethanol, perhaps under defined standard conditions such the boiling point at one atmosphere of pressure. The idea of a true but unknown fixed parameter is less plausible for biological experiments; for example, what does it mean to say *θ* is the effect of a drug on proliferating cells? It depends on the cell line used, whether it is an *in vitro* or *in vivo* experiment, the protocol (concentraion, duration of exposure, route of administration, and so on). The effect may vary between production batches of the compound, or between experimenters, or even between the same experimenter on different occasions. It often doesn’t make sense to assume that a single value of *θ* exists.^21^ And yet, the core of the “reproducibility crisis” lies in researchers inability to obtain similar values for unobservable and nonexistent parameters that, *a priori*, are expected to vary. We should only be concerned, but perhaps not surprised, when results of direct replications differ, because biologists are often unable to control all relevant factors, including the heterogeneity of the sample material. From the predictive perspective, parameters are a means to an end – making predictions – and are not ends in themselves. Even if we know the value of a parameter precisely, we still may be unable to predict well. Suppose we toss a coin 10 million times and obtain 5010000 heads and 4990000 tails. The bias of the coin (its propensity to land heads) is 0.50100 (95% CI: 0.50069 to 0.50131), and the p-value testing whether the bias is significantly different from 0.5 (a fair coin) is 2.54 × 10^*−*10^. Despite this exceedingly small p-value, we are unable to predict if the next toss will be heads any better than chance.

Much of the earlier work on statistical prediction used a Bayesian framework because predictions and their uncertainties could be easily calculated. Stigler even argues that Bayesian inference was originally observation-focused, as Thomas Bayes represented a complete lack of knowledge by reference to a data distribution, and not the distribution of a parameter.^22^ We propose using multilevel models, consistent with previous recommendations for dealing with pseudoreplication, but make them Bayesian, and instead of focusing on unobservable parameters, use predictions from the model to draw conclusions about biological entities of interest. Details of Bayesian inference are not discussed here, but Kruschke and McElreath provide excellent introductions,^23,24^ and Gelman and colleagues cover more advanced topics.^25^ The key idea however is that our uncertainty in all unknowns before seeing the data, such as a parameter in a statistical model, is represented with a distribution, known as the *prior distribution* (Fig. 1). The prior is then updated with the data to form the *posterior distribution*, representing what the data told us about the parameter, combined with what we knew before seeing the data. Predictions are then made from the posterior distribution, giving us the *posterior predictive distribution*, which quantifies our uncertainty in the prediction. A final technical term is the *experimental unit*, which is the biological entity that is randomly and independently assigned to an experimental condition, and it may differ from the biological entity that a scientist wants to make an inference about. The number of experimental units equals the number of genuine replicates, and thus the sample size.^6,7^

**Figure 1:**
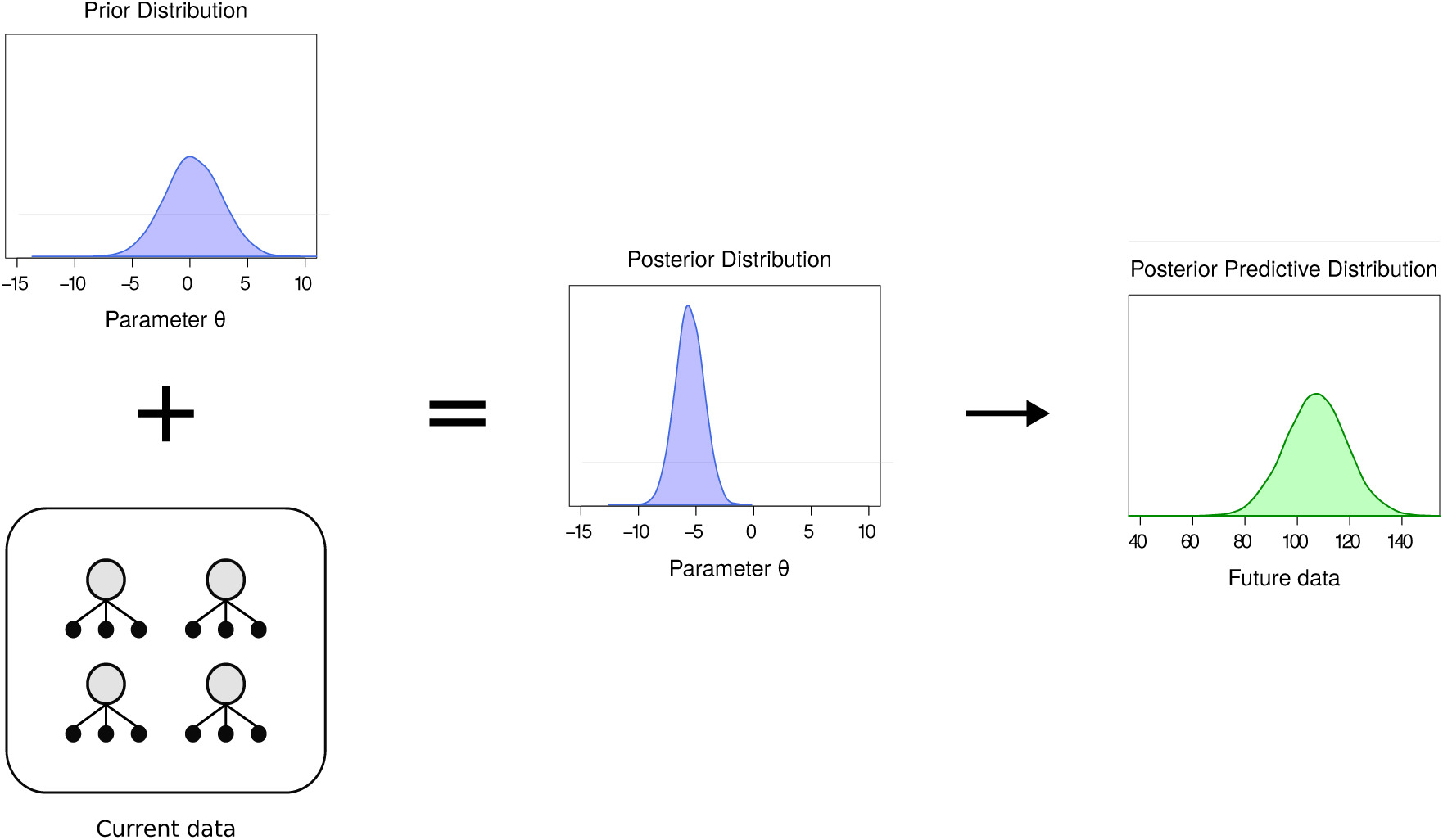
Bayesian updating and predictions. The prior distribution represents what we know about a parameter before seeing the data. The data updates the prior to form the posterior distribution, which is then used to make a prediction about future data through the posterior predictive distribution.

We first analyse a dataset where mice (genuine replicates) were randomly assigned to different conditions and the soma size of multiple neurons (pseudoreplicates) per animal were measured. We compare results from an incorrect analysis, averaging the pseudoreplicates, a classic multilevel model, and a Bayesian multilevel model. Next, we analyse an experiment where pregnant female rodents were randomised to conditions but the interest is in the offspring. Again, we compare the results of several analyses. We also show the effect of doubling the number of replicates versus doubling the number of pseudoreplicates, and how this affects parameter estimates and predictions.

## Methods

### Fatty acid dataset

Moen, Fricano, and colleagues were interested in factors affecting the soma size of neurons *in vivo*.^10,26^ They examined the effect of (1) knocking down the phosphatase and tensin homolog (Pten) gene, (2) infusion of fatty acids (FA) via osmotic minipumps, and (3) the interaction of Pten knockdown and FA infusion on the soma size. Only a subset of the data are used here to highlight the key points with a simpler analysis. In the subset of data, nine mice (genuine replicates) were randomly assigned to a vehicle control or fatty acid condition, and the soma size of 354 neurons (pseudoreplicates) were measured across all animals. Only the control (Pten expressing) neurons are used.

The data were analysed using four methods. The first analysis is the “wrong N” model where all the values are pooled, ignoring the animal-to-animal variability and ignoring pseudoreplication. A two-tailed independent-samples t-test assuming equal variances was used and with an incorrect sample size of 354 neurons. Second, multiple values per animal are averaged and the nine means are analysed with the same t-test, which is an appropriate analysis. Third, a multilevel model appropriately treats neurons as pseudoreplicates by keeping track of which neurons come from which animal. We can write the model as

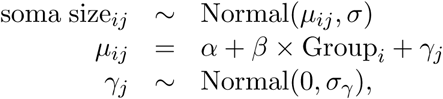

where the soma size for neuron *i* in animal *j* is modelled as coming from a normal distribution that has a mean of *µ* and standard deviation of *σ* (*σ* represents the within-animal variation of soma sizes). *µ* is a deterministic function of the parameters *α*, *β*, and *γ*. *α* is the intercept and represents the mean of the neurons in the control group. *β* is the difference between the vehicle control and FA group and is the key parameter for testing the effect of the treatment. *γ* captures the variation between animals (but within a treatment group) and it is indexed with the subscript *j* because there is one value for each of the nine animals. These nine values are also modelled as coming from a normal distribution with a mean of zero and a standard deviation of *σ*_*γ*_. *α*, *β*, *σ*, and *σ*_*γ*_ are the unknown parameters that we estimate from the data.

The final analysis uses a Bayesian multilevel model that is identical to the model above, but since a Bayesian analysis requires our prior uncertainty about all unknowns to be specified, we use the sensible default priors calculated by the brm() function from the brms R package, which are shown below. The brms() function does not calculate a prior for the difference in group means (*β*); we therefore use a Normal(0, 20) prior, which is wide relative to the size of the expected effect and therefore has little influence on the results.

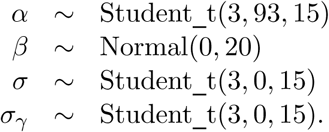

Once the model is fit, we can calculate the posterior predictive distributions by sampling values (neuron sizes) from the model and performing the calculations of interest. You can think of it as sampling an animal from a population of animals, and then sampling a neuron from the population of neurons in that animal. We can then calculate the probability that a randomly sampled neuron from one condition is higher than a randomly sampled neuron from another condition.

### Valproic acid dataset

The data are originally from Mehta and colleagues and a subset of the data from the labstats R package is used here.^6,27,28^ In this smaller dataset, six pregnant dams were randomised to receive either saline (SAL) or valproic acid (VPA) injections. After the offspring were born, four were selected from each litter and randomised to a saline (SAL) or a 2-methyl-6-phenylethylpyrididine (MPEP) drug condition. VPA induces autism-like behaviour in rodents and the aim of the study was to see if the mGluR5-receptor antagonist MPEP improves behaviour. Here, we only examine locomotor activity, although the experiment measured several outcomes. The experiment is a split-plot or split-unit design because there are two types of experimental units and therefore two sample sizes. The first is the six pregnant dams when testing the VPA effect, and the second is the 24 offspring when testing the MPEP effect.

The data were analysed using three methods. The first analysis is the “wrong N” model that ignores the split-unit design of the study and uses a 2-way ANOVA with Group (VPA vs. SAL) and Drug (MPEP vs. SAL) as factors. The test for the drug effect is appropriate, but not for the VPA effect. Second, using a multilevel model we can specify that pregnant dams are the experimental units for the VPA effect and that the offspring are the experimental units for the MPEP effect, thus correctly distinguishing between genuine replication and pseudoreplication. The final model is the Bayesian multilevel model from which we can calculate the predictive distributions. Once again we use the default priors calculated by the brm() function and specify Normal(0, 20) priors for the Drug, Group, and Drug by Group interaction effects.

To better understand how the sample size affects parameter estimates and predictions, we created two new datasets by taking the 24 locomotor activity values and adding random noise from a Normal(0, 0.2) distribution. We then combine these values with the original 24 values to double the number of data points. For one dataset we kept the same number of litters but doubled the number of animals per litter, and in the second we doubled the number of litters. We then compare the results from the original analysis to cases where we have more litters, or the same number of litters but more animals per litter.

All analyses were done with R (version 3.6.1). The Bayesian models were implemented in Stan (version 2.19.2)^29^ via the brms package (version 2.10.0).^30^ The R code to reproduce the figures and results are in Supplementary File 1.

## Results

### Fatty acid dataset

Figure 2A plots all the data, making the sources of between-animal and within-animal variation clear. Figure 2B plots the results as they are commonly seen in journals: groups means and misleading standard errors. This graph is inappropriate because it fails to distinguish between the within-animal and between-animal variability, and because the error bars are based on the incorrect sample size, and are therefore too small. Analysing this data with an inappropriate t-test gives a significant p-value: *t*_(352)_ = −2.27, p = 0.024, N=354. Figure 2C plots the animal averaged values, but the error bars are now based on the correct sample size of nine. The mean of the control group differs in Figure 2B and C because the number of observations differs between animals. Animal 3 has only two soma size measurements and therefore contributes 2/164 = 1.2% of the values when calculating the mean of the control group, but 1/5 = 20% of the values when first calculating mouse means and then using these mean values to calculate the mean of the control group. The error bars for the FA group are similar in Figure 2B and C, despite a difference in sample size. This occurs because there are more values in Figure 2B but they are variable, whereas in Figure 2C there are fewer animal-averaged values but they vary less. The variability and sample sizes happen to balance out such that the error bars are similar, but often the error bars will be considerably larger for the animal-averaged values. The visual impression of an effect differs between Figure 2B and C, and the analysis of the averaged data confirms that there is much less evidence for a group difference: *t*_(7)_ = −1.77, p = 0.12, N=9. Analysing the data with a multilevel model gives similar results as the averaging and t-test method: *t*_(7)_ = −1.49, p = 0.18, N=9. If all animals had the same number of neurons, then averaging and a multilevel model would give identical results (as would a fixed effects nested model that wasn’t discussed).

**Figure 2:**
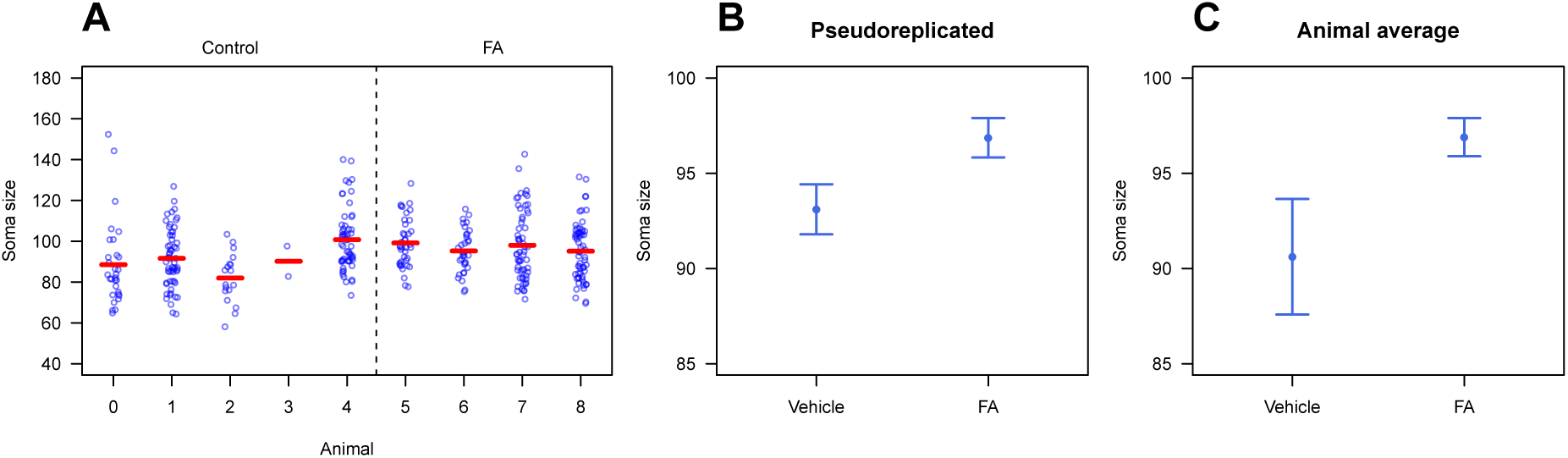
Data from the fatty acid experiment. Plotting the raw values shows the within- and between-animal variation (A). Means and the incorrectly calculated standard errors (B) and correctly calculated standard errors (C).

Figure 3A shows the posterior distribution for the difference between the two groups from the Bayesian analysis. This distribution represents the uncertainty in the difference, given the data, model, and prior information. It focuses on the unobservable group level mean-difference parameter, and is a Bayesian version of the classic multilevel analysis. We can calculate the proportion of the posterior distribution that is below zero, which equals 0.9. The interpretation is that there is a 90% chance that the control group has a lower mean than the fatty acid group. Alternatively, we can calculate the probability in the other tail of the distribution (above zero), and by multiplying this value by two we obtain a value that is often a similar magnitude to a classic p-value. This value is 0.20, and the p-value from the classical multilevel model is 0.18.

**Figure 3:**
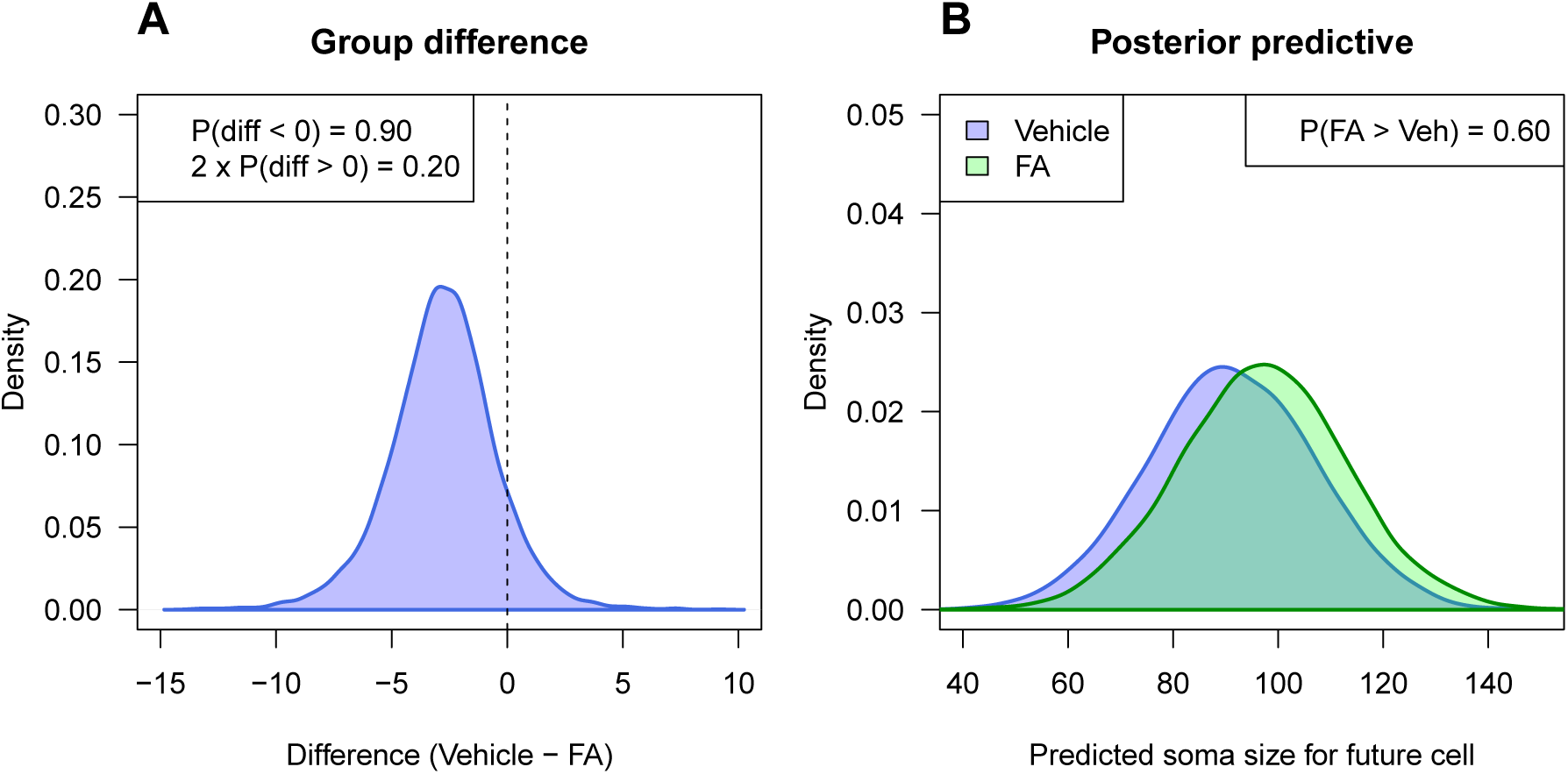
Bayesian results for the fatty acid data. Density plot for the difference between groups (A). The probability that the effect is negative is 0.90 and equals the proportion of the distribution below zero. Two times the proportion of the distribution above zero gives a value similar to a classic two-tailed p-value and equals 0.20 (compare with p=0.18 from the classic multilevel model). Density plot predicting the value of one neuron from a future animal from each experimental condition (B). The probability that randomly selected neuron in the FA condition is higher than a randomly selected neuron in the vehicle control condition is only 0.60.

If we’re interested in making an inference about neurons, we can turn to the predictive perspective and ask: “what is the probability that a randomly chosen neuron from a future animal in the treated group will have a higher value that a randomly chosen neuron from a future animal in the control group?” If the scientific interest is in the individual neurons, then making a probabilistic prediction about as yet unseen neurons directly addresses this question. To obtain these distributions, we first sample an animal from a population of animals, and then sample a neuron within this animal, and then repeat this process thousands times for animals in both experimental conditions. Figure 3B plots these distributions, and they have substantial overlap. The probability that a neuron from the treated group has a larger soma size is only 0.60. Compare this with the p-value of 0.024 incorrectly calculated from the first t-test analysis, which implies evidence for an effect (to the extent that we can use a small p-value to indicate that an effect exists).

A benefit of the predictive approach is that we are making a claim about a future observable event and therefore can test our prediction. 95.3% of the values in the current dataset fall within the 95% prediction interval calculated from the posterior predictive distribution, indicating that we created a suitable model. But the real test of the model is seeing how well we can predict values from treated and control animals when the experiment is run again. Another benefit is that the predictions do not necessarily improve as the number of neurons per animal increase; researchers cannot ramp up the (incorrect) sample size to make small but negligible differences between groups statistically significant.

### Valproic acid experiment

The data from the VPA experiment are plotted in Figure 4 and broken down by Drug, Group, and Litter. Litters C, E, and D are in the saline group and the other litters received VPA prenatally. Black lines connect the group means, and in all litters the MPEP animals have more locomotor activity than the saline animals, on average. The VPA animals also appear to have less activity than animals in the saline group.

**Figure 4:**
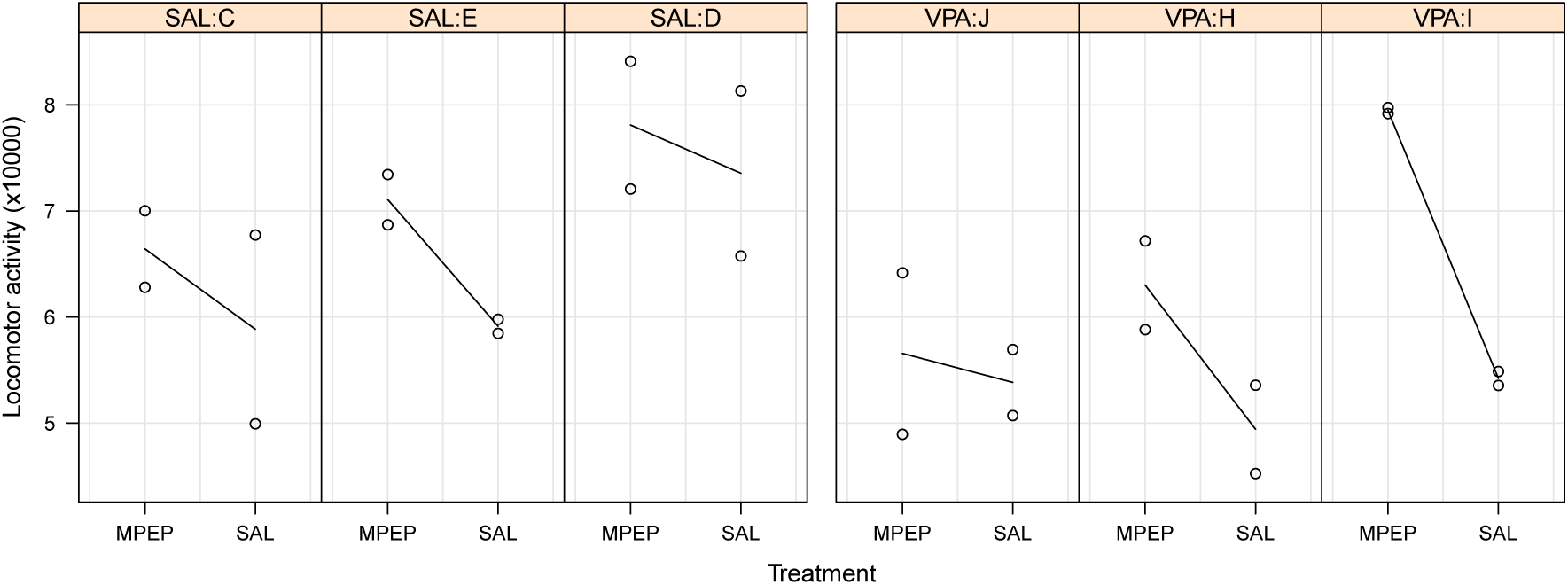
Locomotor activity data for the split-unit valproic acid experiment. Each panel is a litter, which were randomised to receive saline (SAL) or valproic acid (VPA) prenatally. Each point is an offspring, and after birth, animals within a litter were randomised to either a saline control group (SAL) or the drug MPEP.

When these data are analysed with an (inappropriate) 2-way ANOVA the results are: effect of MPEP: F_(1,20)_ = 8.95, p = 0.007, N =24; effect of VPA: F_(1,20)_ = 5.32, p = 0.032, N= 24; and the interaction effect: F_(1,20)_ = 0.64, p = 0.43, N = 24. Thus we would conclude that MPEP increases locomotor activity and VPA decreases it.

Next, we analyse the data with a multilevel model and the results are: MPEP effect: t_(16)_ = 3.62, p=0.002, N=24; VPA effect: t_(4)_ = 1.53, p=0.20, N=6; interaction effect: t_(16)_ = −0.97, p=0.35, N=24. The VPA effect is no longer significant because the multilevel model correctly treats N as the 6 litters and not the 24 offspring. But both analyses give similar results for the MPEP and interaction effects because the sample size is the number of offspring.

Once again, a researcher could argue that they are interested in the effect of VPA on the *offspring*, not on some abstract “litter-averaged” value. We therefore fit a Bayesian version of the multilevel model and obtain similar Bayesian p-values for the effect of MPEP: p = 0.004, VPA: p = 0.24, and the interaction: p = 0.38. The posterior predictive distributions are shown in Figure 5. In Figure 5A we ask: “what is the probability that an animal in a future VPA litter will be less active than an animal in a future control litter, given that both these animals receive saline postnatally?” This question directly focuses on the effect of VPA in the offspring, and the probability is 0.76. This value is less impressive than the incorrect p-value from the 2-way ANOVA of 0.032. Nevertheless, it does suggest that knowing whether an animal received VPA or saline injections prenatally provides some predictive ability, but whether this is large enough is a scientific and not a statistical consideration. The other plots in Figure 5 show predictions for other key comparisons.

**Figure 5:**
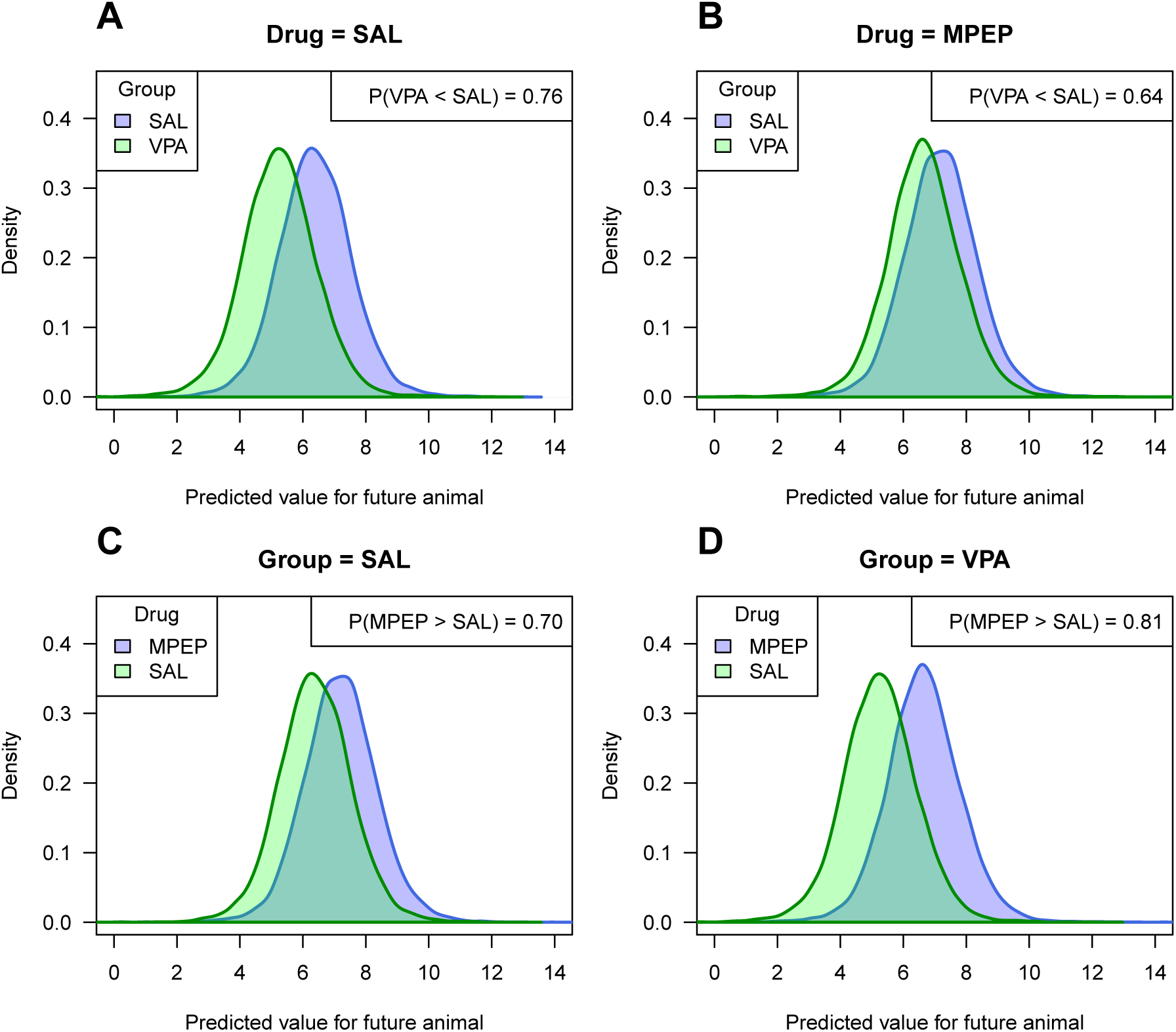
Posterior predictive distributions for the VPA experiment. For animals injected with saline postnatally, the probability is 0.76 that an animal receiving VPA prenatally is less active than an animal receiving saline prenatally (A), and for animals receiving MPEP postnatally, the probability is 0.64 (B). For animals in the prenatal saline litters, the probability is 0.70 that an animal receiving MPEP postnatally is more active than an animal receiving saline (C), and for animals in the prenatal VPA litters, the probability is 0.81 (D).

### Allowing the drug effect to vary by litter

The drug by group interaction tests if the effect of MPEP differs between the VPA and saline groups. Graphically, it tests whether the average slope in the VPA litters differs from the average slope in the saline litters (Fig. 4). Regardless of the result, the effect of MPEP might also differ across litters *within* a treatment group. For example, the difference between MPEP and saline groups is large for litter I, small for litter J, and intermediate for litter H (Fig. 4). The question is: are these differences consistent with sampling variation, or are the effects so different across litters that we suspect that a technical or biological factor is influencing the results?

To address this question, we fit a second Bayesian model (Model 2) to the data where the size of the MPEP drug effect is allowed to vary by litter, and there was little evidence that the size of the MPEP effect differed across litters. The simpler model without the interaction is 18 times more likely compared with Model 2 when comparing the models with a Bayes factor.^31^ This new source of variation increases the uncertainty in the predictions because observed values can now span a wider range. The probabilities calculated in Figure 5 are shown again in Figure 6 for both models. The probabilities are closer to random guessing (0.5) for Model 2 because another source of uncertainty has been included in the predictions, and when we calculate the probability that a randomly selected animal in the VPA condition is less active than a randomly selected animal in the saline condition, we now include the additional variability across the litters. Since there was little evidence for a drug by litter interaction, the predictions from Model 2 differ only slightly from those of Model 1, but it nevertheless illustrates how different sources of uncertainty can be incorporated into the predictions, and how it affects the predicted values.

**Figure 6:**
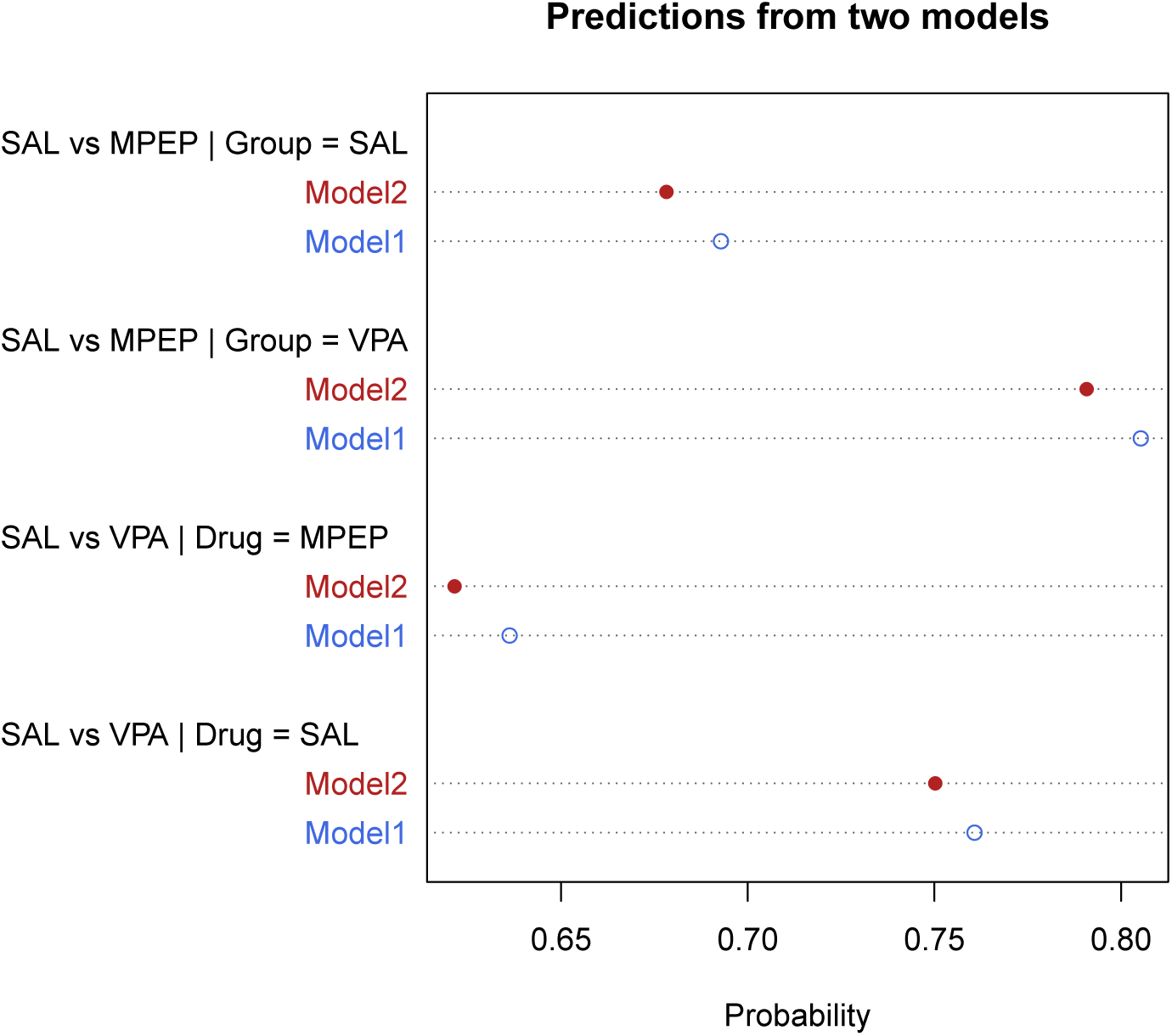
Prediction probabilities for two models. Model 1 assumes that MPEP’s effect is constant across litters while Model 2 allows for the effect to be larger in some litters and smaller in others. Since Model 2 includes another source of uncertainty, the predictions are more uncertain and therefore the calculated probabilities are closer to random guessing (0.5).

### Effect of doubling the number of litters versus doubling the number of animals per litter

To better understand how the sample size affects parameter estimates versus predictions, we created two new datasets. For the first dataset we kept the same number of litters but doubled the number of animals per litter; in the second, we doubled the number of litters. Thus we can compare the results from the original analysis to cases where we have more litters, or the same number of litters but more animals per litter. The same Bayesian model was fit to these new datasets and the results are shown in Figure 7. Doubling the number of litters or the number of animals per litter increases the precision of the MPEP effect (the posterior distribution is narrower, and this would make a classic p-value smaller; Fig. 7A). The posterior distribution for the original VPA effect (Fig. 7B, blue distribution, although it is obscured by the red distribution) is wider than the original MPEP effect in Figure 7A, indicating that the result is less certain, which we expect because N=6 for testing the VPA effect and N=24 for testing the MPEP effect. Doubling the number of litters increases the precision of the VPA effect (Fig. 7B; green distribution), but not doubling the number of animals per litter (red distribution). This result is expected and is why the number of genuine replicates (litters) must be increased to improve power to detect a VPA effect.

**Figure 7:**
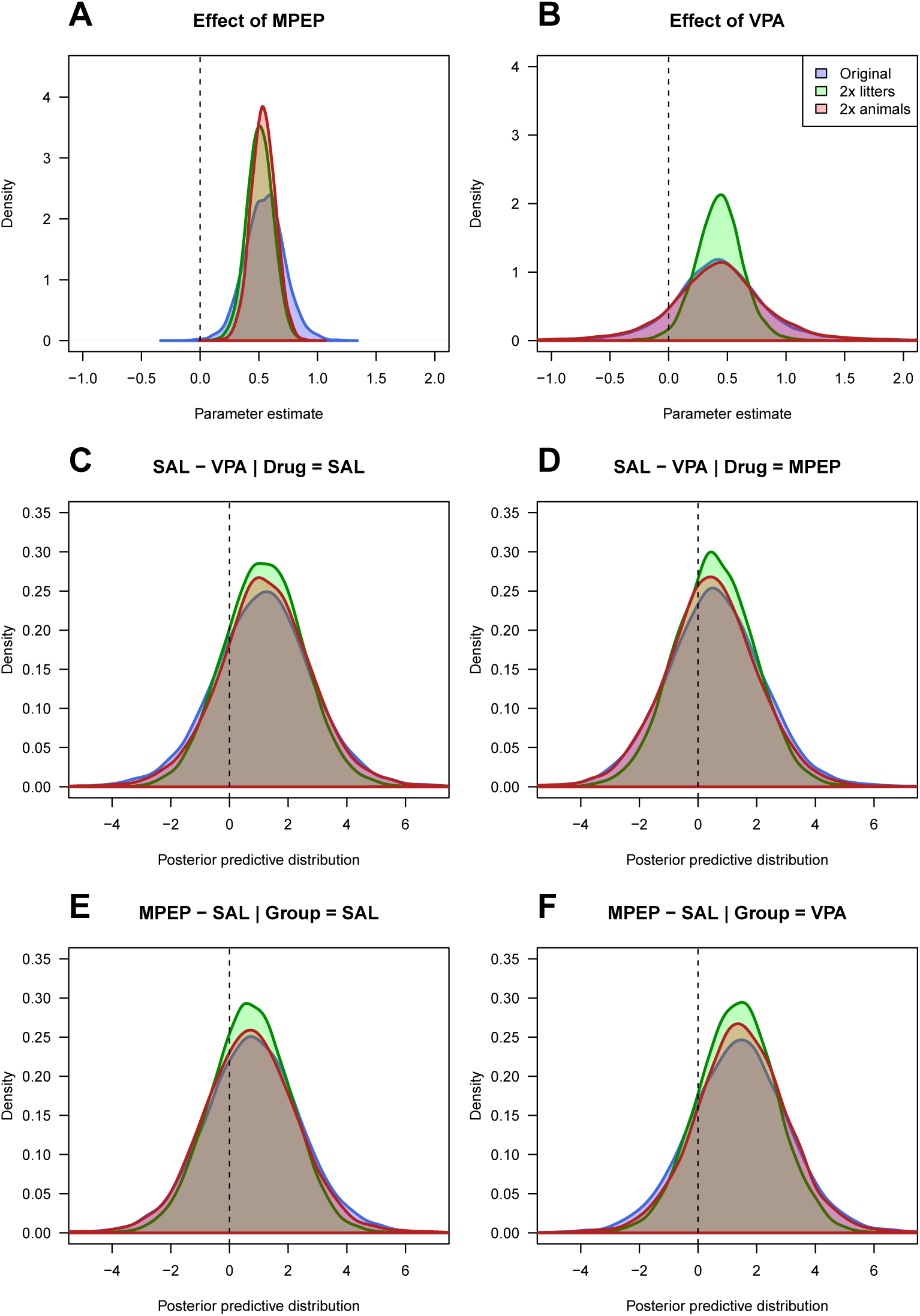
Effect of doubling the number of litters versus doubling the number of animals per litter. Panel (A) and (B) show the effect on the parameter estimates. For MPEP, the sample size is the number of offspring, and so precision is increased (narrower distributions) in both cases. For VPA, the sample size is the number of litters, and only increasing the number of litters increases the precision of this estimate. Panels (C–D) compare the posterior predictive distributions, which get only slightly narrower.

The blue distributions in Figures 7C–F plot the same results as Figure 5A–D, but instead of plotting two distributions, the distribution of their difference is plotted, which is a more compact representation. The other two distributions plot the same information from the datasets that doubled the number of litters (green) or the number animals per litter (red), which have only slight increases in the precision of the posterior predictive distributions, again, reflecting that further knowledge of a parameter does not necessarily lead to better predictions.

## Discussion

An old statistical framework is being revived to improve reproducibility in science by focusing on predicting observable outcomes and downgrading the importance of testing hypothetical parameters. The predictive approach also solves a problem related to pseudoreplication: inferences can be made about entities of direct interest to biologists. We made probabilistic conclusions about neurons in the first example and about offspring in the second example, both of which are pseudoreplicates.

Graphs like Figure 2B can mislead scientists into thinking that the effect is biologically relevant, given the distance between means, relative to the length of the error bars. The incorrectly calculated p-value of 0.024 further reinforces this conclusion. We note the incoherence of arguing that the scientific interest is in the individual neurons, but then presenting only means and error bars where information about the neurons is hidden, and the neuron-to-neuron variation is mixed up with the animal-to-animal variation. Graphs such as Figure 2A should be presented, if not in the main manuscript, then at least in the supplementary material. Figure 2A shows the variability of values within animals, making it obvious that the observed mean difference between groups of 2.8 µm^2^ is small compared with the range of values within an animal, 95% of which will be between 84.4 and 115.8 µm^2^. The predictive approach provides an alternative way to interpret results, although it still may be difficult to understand the biological significance of these results. However, an analogy can be made with a clinical trial. If you are considering a medical treatment, you want to know the probability that you will be better off under treatment A versus treatment B, which can be calculated from the posterior predictive distribution.^17^ A p-value and a population-averaged effect size do not provide this information, and with a large sample size, the clinical trial could return a highly significant p-value, but the probability of a better outcome under treatment A could be close to a coin flip. Demidenko makes a similar point from a frequentist perspective.^32^ Similarly, if large soma sizes are “good”, we can calculate the probability that our experimental intervention will give us this desired outcome for a single cell, or for any numbers cells (which we did not calculate, but it is straightforward). Calculating such summaries from the posterior predictive distributions can therefore complement the usual parametric results.

The approach described here can be applied to all data types and models, and is easily extendable. For example, we could have fit a model that had different variances for the control and fatty acid groups in the first example, or we could allow each animal to have a different variance for the distribution of soma sizes. The calculations for the predictions are identical to the current examples. Sometimes two or more models are equally good and at describing the data and it is unclear which should be used for predictions. Fortunately, we do not need to choose only one model but can make predictions from several, and then combine the results and weighting them according to how well the models fit the data.^33^ Here, we would also include model uncertainty when making predictions.

The predictive approach could also be beneficial when experiments are conducted in batches (e.g. see Karp and colleagues, this issue) or if data accumulate over time. Here we can test our ability to predict ahead to data that we will obtain, and thus be able to test our predictions on future observations.

Poor predictive ability has direct consequences for interpreting preclinical animal research. If we cannot predict well in the animal models that we use as proxies for human diseases, what hope is there for using animals to predict what will happen in humans?

## Acknowledgements

JRM’s research is funded by the Wellcome Trust and Biological Sciences Research Council (BBSRC). MCA’s research is funded by the Medical Research Council (MR/J013188/1), EUFP17 Marie Curie Actions (PCIG10-GA-2011-303680), and the BBSRC (BB/R002177/1). MRM’s research is funded by the Medical Research Council (MC_UU_00011/7).

## Author contributions

SL: Conceived and designed the analysis, performed the analysis, wrote the paper. JM, MA, MM, helped define the problem, discussed the results, provided critical feedback, and contributed to the final manuscript.

## Data availability

The data used are already publicly available, but have been included in the Supplementary File 1 for convenience.

## Supplementary Information

Supplementary File 1. Data_and_code.zip contains the data and R code.

