## Supplementary material for "A Bayesian predictive approach for dealing with pseudoreplication": Data and R code: code.R

```
library(tidyverse)
library(labstats) # contains the VPA data
library(brms)
library(bayesplot)
library(nlme)
library(sciplot)

options(mc.cores = parallel::detectCores())

## use sum-to-zero contrasts
options(contrasts=c("contr.sum", "contr.poly"))


## ----------------------------------------------------------------------
## Moen data
## ----------------------------------------------------------------------


## read data
dat <- read.csv("Moen_data.csv")


## select key groups, variables, and recode factor
dat %>% filter(pten == 0 & fa != 2) %>%
    select(mouseid, fa, somasize) %>%
    mutate(fa=recode_factor(fa, `0`="Vehicle", `1`="FA"),
           mouseid=factor(mouseid)) ->
    d


## calculate animal average for alternate analysis
d %>%
    group_by(mouseid, fa) %>%
    summarise(y.mean=mean(somasize)) %>%
    data.frame() ->
    d.red

pdf("Fig2.pdf", height=3, width=10)
par(las=1,
    mfrow=c(1, 3),
    mar = c(4.5, 4, 3.5, 1),
    cex=2)

layout(matrix(c(1, 2, 3), nrow=1),
       width=c( 0.4, 0.3, 0.3),
       height=1)
stripchart(somasize ~ mouseid, data=d, vertical = TRUE, method="jitter", pch=1,
           ylab="Soma size", xlab="Animal", ylim=c(40, 180),
           col=rgb(0, 0, 1, 0.5))
abline(v=5.5, lty=2)
par(xpd=TRUE)
text(x=c(3, 7.5), y=c(195, 195), label=c("Control", "FA"))
par(xpd=FALSE)

segments(c(1:9) - 0.2, d.red$y.mean, c(1:9) + 0.2, d.red$y.mean,
         lwd=3, col="red")
mtext("A", side=3, line=1.5, adj=0, font=2, cex=1.5)

lineplot.CI(fa, somasize, data=d, type="p", ylim=c(85, 100), xlim=c(0.5, 2.5),
            xlab="", ylab="Soma size", col="royalblue", lwd=1.5,
            main="Pseudoreplicated")
mtext("B", side=3, line=1.5, adj=0, font=2, cex=1.5)

lineplot.CI(fa, y.mean, data=d.red, type="p", ylim=c(85, 100), xlim=c(0.5, 2.5),
            xlab="", ylab="Soma size", col="royalblue", lwd=1.5,
            main="Animal average")
mtext("C", side=3, line=1.5, adj=0, font=2, cex=1.5)

dev.off()


## Incorrect analysis of all values with t-test
t.test(somasize ~ fa, data=d, var.equal = TRUE)

## Correct analysis of animal-averagted values with t-test
t.test(y.mean ~ fa, data=d.red, var.equal = TRUE)

## Correct analysis of all values by specifying the error structure
## (not discussed in paper but here for a comparison)
summary(aov(somasize ~ fa + Error(mouseid), data=d))

## Correct analysis of all values with a hierarchical model
summary(lme(somasize ~ fa, random= ~ 1 | mouseid, data=d))


## Proposed Bayesian analysis of all data
## examine default priors
get_prior(bf(somasize ~ fa + (1 | mouseid), center = TRUE), data=d)

## put prior on parameter for treatment effect
m1.prior <- c(
    prior(normal(0, 20), class="b", coef="fa1")
)

## fit model
m1 <- brm(somasize ~ fa + (1 | mouseid), data=d,
          iter = 10000, chains = 3, seed=123,
          prior=m1.prior, 
          control=list(adapt_delta=0.99))


## posterior predictive checks
pp_check(m1, nsamples = 100)
pp_check(m1, type="stat")
pp_check(m1, type="stat", stat="sd")
pp_check(m1, type="stat", stat="max")
pp_check(m1, type="stat", stat="min")
pp_check(m1, type="intervals")

## extract parameter for the difference in group means
post_diff <- posterior_samples(m1)$b_fa1

## probability that effect is negative
mean(post_diff < 0)

## calculate a value similar to a classic p-value
2 * mean(post_diff > 0)

## new data used to make predictions
new.data <- data.frame(mouseid=c(NA, NA), fa=c("Vehicle", "FA"))


## make predictions for new cells from new animals
preds <- posterior_predict(m1,
                           newdata = new.data, re_formula =  ~ (1 | mouseid),
                           allow_new_levels=TRUE)


## probability that randomly selected FA cell is higher than a
## randomly selected vehicle cell
mean(sample(preds[, 2]) > preds[, 1])


## how many cells fall within prediction interval for new animal in each group
p1 <- mean(d$somasize[d$fa == "Vehicle"] > quantile(preds[, 1], 0.025) &
           d$somasize[d$fa == "Vehicle"] < quantile(preds[, 1], 0.975))

p2 <- mean(d$somasize[d$fa == "FA"] > quantile(preds[, 2], 0.025) &
           d$somasize[d$fa == "FA"] < quantile(preds[, 2], 0.975))

(p1 + p2)/2


## plots of results from Bayesian model
pdf("Fig3.pdf", height=4.5, width=9)
par(las=1,
    mfrow=c(1, 2),
    mar = c(4.5, 4, 3, 1),
    cex=1)

## ----- 1
dens <- density(post_diff, adjust=1.25)
plot(dens, main="Group difference",
     xlab="Difference (Vehicle - FA)", xlim=c(-15, 10), ylim=c(0, 0.3))

polygon(c(-0.01, dens$x, 1.01), c(0, dens$y, 0),
        col=rgb(0, 0, 1, 0.25), border="royalblue", lwd=2)

abline(v= 0, lty=2)
mtext("A", side=3, line=1.5, adj=0, font=2, cex=1.5)
legend("topleft", legend = c("P(diff < 0) = 0.90",
                              "2 x P(diff > 0) = 0.20"))


## ----- 2
dens1 <- density(preds[, 1], adjust=1.25)
dens2 <- density(preds[, 2], adjust=1.25)

plot(dens1, xlab="Predicted soma size for future cell",
     main="Posterior predictive",
     ylim = c(0, 0.05), xlim = c(40, 150))
polygon(c(-0.01, dens1$x, 1.01), c(0, dens1$y, 0),
        col=rgb(0, 0, 1, 0.25), border="royalblue", lwd=2)

polygon(c(-0.01, dens2$x, 1.01), c(0, dens2$y, 0),
        col=rgb(0, 1, 0, 0.25), border="green4", lwd=2)

mtext("B", side=3, line=1.5, adj=0, font=2, cex=1.5)
legend("topleft", col=c("royalblue", "green4"),
       fill=c(rgb(0, 0, 1, 0.25), rgb(0, 1, 0, 0.25)), 
       legend = c("Vehicle", "FA"))
legend("topright", legend = "P(FA > Veh) = 0.60")

dev.off()


## ----------------------------------------------------------------------
## Valproic acid data (from labstats package)
## ----------------------------------------------------------------------

## scale activity values for nicer plots
VPA %>%
    mutate(activity=activity / 10000) ->
    VPA


pdf("Fig4.pdf", height=4.5, width=11)
lattice::trellis.par.set(axis.components=list(right=list(tck=0),
                                              top=list(tck=0)))

lattice::xyplot(activity ~ drug|group:reorder(litter, activity, mean),
                data=VPA, type=c("g","p","a"), col="black", layout=c(6,1),
                xlab="Treatment", ylab="Locomotor activity (x10000)",
                scales=list(alternating=FALSE), between=list(x=c(0,0,1,0,0)))
dev.off()


## Incorrect analysis
car::Anova(lm(activity ~ drug * group, data=VPA))

## Correct analysis by specifying the error structure
## (not discussed in paper, but here for a comparison)
summary(aov(activity ~ drug * group + Error(litter), data=VPA))

## Correct analysis of all values with a hierarchical model
summary(lme(activity ~ drug * group, random= ~ 1 | litter, data=VPA))


## Proposed Bayesian analyses
## Examine default priors
get_prior(activity ~ drug * group + (1 | litter), data=VPA)

## put prior on parameters for treatment effects
m2.prior <- c(
    prior(normal(0, 20), class="b")
)

## fit model
m2 <- brm(activity ~ drug * group + (1 | litter), data=VPA,
          iter = 10000, chains = 3, seed=123,
          prior=m2.prior, save_all_pars=TRUE, 
          control=list(adapt_delta=0.999))

## A second model where we allow the size of the drug effect to vary by litter
m2b <- brm(activity ~ drug * group + (drug | litter), data=VPA,
           iter = 10000, chains = 3, seed=123,
           prior=m2.prior, save_all_pars=TRUE,
           control=list(adapt_delta=0.999))


## new data used to make predictions
new.vpa <- VPA[!duplicated(VPA[, 2:3]), 2:3]
new.vpa$litter <- c("P", "P", "Q", "Q")
new.vpa$litter <- rep(NA, 4)

## make predictions for new offspring from new dams
m2.preds <- posterior_predict(m2,
                              newdata = new.vpa, re_formula = ~ 1 | litter,
                              allow_new_levels=TRUE)


m2b.preds <- posterior_predict(m2b,
                              newdata = new.vpa, re_formula = ~ drug | litter,
                              allow_new_levels=TRUE)


pdf("Fig5.pdf", height=7, width=8)
par(las=1,
    mfrow=c(2, 2),
    mar = c(4.5, 4, 3, 1))

## ----- 1
dens1 <- density(m2.preds[, 2], adjust=1.25)
dens2 <- density(m2.preds[, 4], adjust=1.25)

plot(dens1, xlab="Predicted value for future animal",
     main="Drug = SAL",
     ylim = c(0, 0.45), xlim = c(0, 14))
polygon(c(-0.01, dens1$x, 1.01), c(0, dens1$y, 0),
        col=rgb(0, 0, 1, 0.25), border="royalblue", lwd=2)

lines(dens2)
polygon(c(-0.01, dens2$x, 1.01), c(0, dens2$y, 0),
        col=rgb(0, 1, 0, 0.25), border="green4", lwd=2)

mtext("A", side=3, line=1.5, adj=0, font=2, cex=1.5)
legend("topleft", col=c("royalblue", "green4"),
       fill=c(rgb(0, 0, 1, 0.25), rgb(0, 1, 0, 0.25)), 
       legend = c("SAL", "VPA"), title="Group")

legend("topright", legend="P(VPA < SAL) = 0.76")


## ----- 2
dens1 <- density(m2.preds[, 1], adjust=1.25)
dens2 <- density(m2.preds[, 3], adjust=1.25)

plot(dens1, xlab="Predicted value for future animal",
     main="Drug = MPEP",
     ylim = c(0, 0.45), xlim = c(0, 14))
polygon(c(-0.01, dens1$x, 1.01), c(0, dens1$y, 0),
        col=rgb(0, 0, 1, 0.25), border="royalblue", lwd=2)

lines(dens2)
polygon(c(-0.01, dens2$x, 1.01), c(0, dens2$y, 0),
        col=rgb(0, 1, 0, 0.25), border="green4", lwd=2)

mtext("B", side=3, line=1.5, adj=0, font=2, cex=1.5)
legend("topleft", col=c("royalblue", "green4"),
       fill=c(rgb(0, 0, 1, 0.25), rgb(0, 1, 0, 0.25)), 
       legend = c("SAL", "VPA"), title="Group")
legend("topright", legend="P(VPA < SAL) = 0.64")


## ----- 3
dens1 <- density(m2.preds[, 1], adjust=1.25)
dens2 <- density(m2.preds[, 2], adjust=1.25)

plot(dens1, xlab="Predicted value for future animal",
     main="Group = SAL",
     ylim = c(0, 0.45), xlim = c(0, 14))
polygon(c(-0.01, dens1$x, 1.01), c(0, dens1$y, 0),
        col=rgb(0, 0, 1, 0.25), border="royalblue", lwd=2)

lines(dens2)
polygon(c(-0.01, dens2$x, 1.01), c(0, dens2$y, 0),
        col=rgb(0, 1, 0, 0.25), border="green4", lwd=2)

mtext("C", side=3, line=1.5, adj=0, font=2, cex=1.5)
legend("topleft", col=c("royalblue", "green4"),
       fill=c(rgb(0, 0, 1, 0.25), rgb(0, 1, 0, 0.25)), 
       legend = c("MPEP", "SAL"), title="Drug")

legend("topright", legend="P(MPEP > SAL) = 0.70")


## ----- 4
dens1 <- density(m2.preds[, 3], adjust=1.25)
dens2 <- density(m2.preds[, 4], adjust=1.25)

plot(dens1, xlab="Predicted value for future animal",
     main="Group = VPA",
     ylim = c(0, 0.45), xlim = c(0, 14))
polygon(c(-0.01, dens1$x, 1.01), c(0, dens1$y, 0),
        col=rgb(0, 0, 1, 0.25), border="royalblue", lwd=2)

lines(dens2)
polygon(c(-0.01, dens2$x, 1.01), c(0, dens2$y, 0),
        col=rgb(0, 1, 0, 0.25), border="green4", lwd=2)

mtext("D", side=3, line=1.5, adj=0, font=2, cex=1.5)
legend("topleft", col=c("royalblue", "green4"),
       fill=c(rgb(0, 0, 1, 0.25), rgb(0, 1, 0, 0.25)), 
       legend = c("MPEP", "SAL"), title="Drug")

legend("topright", legend="P(MPEP > SAL) = 0.81")

dev.off()


## compare both models and calculate Bayes Factor
bayes_factor(m2, m2b)

## extract samples of parameter estimates
m2.post <- posterior_samples(m2)

## calculate relevant values
2 * mean(m2.post$b_drug1 < 0)
2 * mean(m2.post$b_group1 < 0)
2 * mean(m2.post[, "b_drug1:group1"] > 0)

## ---------------------------------------------
## SAL vs VPA \ Drug = SAL, random litter
a <- mean(sample(m2.preds[, 2]) > m2.preds[, 4])
b <- mean(sample(m2b.preds[, 2]) > m2b.preds[, 4])


## ---------------------------------------------
## SAL vs VPA \ Drug = MPEP, random litter
c <- mean(sample(m2.preds[, 1]) > m2.preds[, 3])
d <- mean(sample(m2b.preds[, 1]) > m2b.preds[, 3])


## ---------------------------------------------
## SAL vs MPEP \ Group = SAL, random litter
e <- mean(sample(m2.preds[, 1]) > m2.preds[, 2])
f <- mean(sample(m2b.preds[, 1]) > m2b.preds[, 2])


## ---------------------------------------------
## SAL vs MPEP \ Group = VPA, random litter
g <- mean(sample(m2.preds[, 3]) > m2.preds[, 4])
h <- mean(sample(m2b.preds[, 3]) > m2b.preds[, 4])


## ---------------------------------------------
## Interaction
mean(sample(m2.preds[, 2]) - m2.preds[, 4] >
     sample(m2.preds[, 1]) - m2.preds[, 3])

mean(sample(m2.preds[, 1]) - m2.preds[, 2] < 
     sample(m2.preds[, 3]) - m2.preds[, 4])


## define plot attributes
cols  <-  rep(c("royalblue", "firebrick"), 4)
pchs <- rep(c(1, 16), 4)
mods <- rep(c("Model1", "Model2"), 4)
comps <- rep(c("SAL vs VPA | Drug = SAL",
           "SAL vs VPA | Drug = MPEP",
           "SAL vs MPEP | Group = SAL",
           "SAL vs MPEP | Group = VPA"), each=2)


pdf("Fig6.pdf", height=6, width=7)

dotchart(c(a, b, c, d, e, f, g, h), 
         xlab="Probability", col=cols, pch=pchs,
         groups=factor(comps), labels=mods, lcolor="grey40",
         main="Predictions from two models")

dev.off()


## simulate data for a new litter
VPA2 <- VPA

VPA2 %>%
    mutate(litter=factor(paste0(litter, 2)),
           activity=activity + rnorm(n(), 0, 0.2)) -> 
    VPA2

VPA.more.litters <- rbind(VPA, VPA2)

m3 <- brm(activity ~ drug * group + (1 | litter), data=VPA.more.litters,
          iter = 10000, chains = 3, seed=123,
          prior=m2.prior, save_all_pars=TRUE, 
          control=list(adapt_delta=0.999))

m3.preds <- posterior_predict(m3,
                              newdata = new.vpa, re_formula = ~ 1 | litter,
                              allow_new_levels=TRUE)

## extract samples of parameter estimates
m3.post <- posterior_samples(m3)

## calculate relevant values
2 * mean(m3.post$b_drug1 < 0)
2 * mean(m3.post$b_group1 < 0)
2 * mean(m3.post[, "b_drug1:group1"] > 0)


## simulate data for more cells in the same litters
VPA3 <- VPA
VPA3 %>%
    mutate(activity=activity + rnorm(n(), 0, 0.2)) -> 
    VPA3

VPA.more.cells <- rbind(VPA, VPA3)


m4 <- brm(activity ~ drug * group + (1 | litter), data=VPA.more.cells,
          iter = 10000, chains = 3, seed=123,
          prior=m2.prior, save_all_pars=TRUE, 
          control=list(adapt_delta=0.999, max_treedepth=12))

m4.preds <- posterior_predict(m4,
                              newdata = new.vpa, re_formula = ~ 1 | litter,
                              allow_new_levels=TRUE)

## extract samples of parameter estimates
m4.post <- posterior_samples(m4)

## calculate relevant values
2 * mean(m4.post$b_drug1 < 0)
2 * mean(m4.post$b_group1 < 0)
2 * mean(m4.post[, "b_drug1:group1"] > 0)


pdf("Fig7.pdf", height=10, width=7)
par(las=1,
    mfrow=c(3, 2),
    mar = c(4.5, 4, 4, 1))

dens1 <- density(m2.post$b_drug1, adjust=1.25)
dens2 <- density(m3.post$b_drug1, adjust=1.25)
dens3 <- density(m4.post$b_drug1, adjust=1.25)

plot(dens1, ylim=c(0, 4.5), xlim=c(-1, 2), 
     main="Effect of MPEP", xlab="Parameter estimate")
polygon(c(-0.01, dens1$x, 1.01), c(0, dens1$y, 0),
        col=rgb(0, 0, 1, 0.25), border="royalblue", lwd=2)

lines(dens2) # more litters
polygon(c(-0.01, dens2$x, 1.01), c(0, dens2$y, 0),
        col=rgb(0, 1, 0, 0.25), border="green4", lwd=2)

lines(dens3) # more animals
polygon(c(-0.01, dens3$x, 1.01), c(0, dens3$y, 0),
        col=rgb(1, 0, 0, 0.25), border="firebrick", lwd=2)

abline(v=0, lty=2)
mtext("A", side=3, line=1.5, adj=0, font=2, cex=1.5)


## ---------
dens1 <- density(m2.post$b_group1, adjust=1.25)
dens2 <- density(m3.post$b_group1, adjust=1.25)
dens3 <- density(m4.post$b_group1, adjust=1.25)

plot(dens1, ylim=c(0, 4), xlim=c(-1, 2),
     main="Effect of VPA", xlab="Parameter estimate")
polygon(c(-0.01, dens1$x, 1.01), c(0, dens1$y, 0),
        col=rgb(0, 0, 1, 0.25), border="royalblue", lwd=2)

lines(dens2, col="royalblue", lwd=2, lty=2) # more litters
polygon(c(-0.01, dens2$x, 1.01), c(0, dens2$y, 0),
        col=rgb(0, 1, 0, 0.25), border="green4", lwd=2)

lines(dens3, col="firebrick", lwd=2, lty=4) # more animals
polygon(c(-0.01, dens3$x, 1.01), c(0, dens3$y, 0),
        col=rgb(1, 0, 0, 0.25), border="firebrick", lwd=2)

abline(v=0, lty=2)
mtext("B", side=3, line=1.5, adj=0, font=2, cex=1.5)

legend("topright", col=c("royalblue", "green4", "firebrick"),
       fill=c(rgb(0, 0, 1, 0.25), rgb(0, 1, 0, 0.25), rgb(1, 0, 0, 0.25)), 
       legend=c("Original", "2x litters", "2x animals"))


## ---------------------------------------------
## SAL vs VPA \ Drug = SAL, random litter
dens1 <- density(sample(m2.preds[, 2]) - m2.preds[, 4], adjust=1.25)
dens2 <- density(sample(m3.preds[, 2]) - m3.preds[, 4], adjust=1.25)
dens3 <- density(sample(m4.preds[, 2]) - m4.preds[, 4], adjust=1.25)

plot(dens1, ylim=c(0, 0.35), xlim=c(-5, 7),
     main="SAL - VPA | Drug = SAL", xlab="Posterior predictive distribution")
polygon(c(-0.01, dens1$x, 1.01), c(0, dens1$y, 0),
        col=rgb(0, 0, 1, 0.25), border="royalblue", lwd=2)

lines(dens2)
polygon(c(-0.01, dens2$x, 1.01), c(0, dens2$y, 0),
        col=rgb(0, 1, 0, 0.25), border="green4", lwd=2)

lines(dens3)
polygon(c(-0.01, dens3$x, 1.01), c(0, dens3$y, 0),
        col=rgb(1, 0, 0, 0.25), border="firebrick", lwd=2)

abline(v=0, lty=2)
mtext("C", side=3, line=1.5, adj=0, font=2, cex=1.5)

## ---------------------------------------------
## SAL vs VPA \ Drug = MPEP, random litter
dens1 <- density(sample(m2.preds[, 1]) - m2.preds[, 3], adjust=1.25)
dens2 <- density(sample(m3.preds[, 1]) - m3.preds[, 3], adjust=1.25)
dens3 <- density(sample(m4.preds[, 1]) - m4.preds[, 3], adjust=1.25)

plot(dens1, ylim=c(0, 0.35), xlim=c(-5, 7),
     main="SAL - VPA | Drug = MPEP", xlab="Posterior predictive distribution")
polygon(c(-0.01, dens1$x, 1.01), c(0, dens1$y, 0),
        col=rgb(0, 0, 1, 0.25), border="royalblue", lwd=2)

lines(dens2, col="royalblue", lwd=2, lty=2)
polygon(c(-0.01, dens2$x, 1.01), c(0, dens2$y, 0),
        col=rgb(0, 1, 0, 0.25), border="green4", lwd=2)

lines(dens3, col="firebrick", lwd=2, lty=4)
polygon(c(-0.01, dens3$x, 1.01), c(0, dens3$y, 0),
        col=rgb(1, 0, 0, 0.25), border="firebrick", lwd=2)

abline(v=0, lty=2)
mtext("D", side=3, line=1.5, adj=0, font=2, cex=1.5)

## ---------------------------------------------
## SAL vs MPEP \ Group = SAL, random litter
dens1 <- density(sample(m2.preds[, 1]) - m2.preds[, 2], adjust=1.25)
dens2 <- density(sample(m3.preds[, 1]) - m3.preds[, 2], adjust=1.25)
dens3 <- density(sample(m4.preds[, 1]) - m4.preds[, 2], adjust=1.25)

plot(dens1, ylim=c(0, 0.35), xlim=c(-5, 7),
     main="MPEP - SAL | Group = SAL", xlab="Posterior predictive distribution")
polygon(c(-0.01, dens1$x, 1.01), c(0, dens1$y, 0),
        col=rgb(0, 0, 1, 0.25), border="royalblue", lwd=2)

lines(dens2, col="royalblue", lwd=2, lty=2)
polygon(c(-0.01, dens2$x, 1.01), c(0, dens2$y, 0),
        col=rgb(0, 1, 0, 0.25), border="green4", lwd=2)

lines(dens3, col="firebrick", lwd=2, lty=4)
polygon(c(-0.01, dens3$x, 1.01), c(0, dens3$y, 0),
        col=rgb(1, 0, 0, 0.25), border="firebrick", lwd=2)

abline(v=0, lty=2)
mtext("E", side=3, line=1.5, adj=0, font=2, cex=1.5)

## ---------------------------------------------
## SAL vs MPEP \ Group = VPA, random litter
dens1 <- density(sample(m2.preds[, 3]) - m2.preds[, 4], adjust=1.25)
dens2 <- density(sample(m3.preds[, 3]) - m3.preds[, 4], adjust=1.25)
dens3 <- density(sample(m4.preds[, 3]) - m4.preds[, 4], adjust=1.25)

plot(dens1, ylim=c(0, 0.35), xlim=c(-5, 7),
     main="MPEP - SAL | Group = VPA", xlab="Posterior predictive distribution")
polygon(c(-0.01, dens1$x, 1.01), c(0, dens1$y, 0),
        col=rgb(0, 0, 1, 0.25), border="royalblue", lwd=2)

lines(dens2, col="royalblue", lwd=2, lty=2)
polygon(c(-0.01, dens2$x, 1.01), c(0, dens2$y, 0),
        col=rgb(0, 1, 0, 0.25), border="green4", lwd=2)

lines(dens3, col="firebrick", lwd=2, lty=4)
polygon(c(-0.01, dens3$x, 1.01), c(0, dens3$y, 0),
        col=rgb(1, 0, 0, 0.25), border="firebrick", lwd=2)

abline(v=0, lty=2)
mtext("F", side=3, line=1.5, adj=0, font=2, cex=1.5)

dev.off()
```
